# Comparison of extracellular vesicles and mechanically induced vesicles for structure determination of membrane proteins

**DOI:** 10.64898/2026.03.04.709473

**Authors:** Chunyang Wang, Ole Østergaard, Roberto Melero, Gabriela Nagy-Davidescu, Matthias Eibauer, Jesper Velgaard Olsen, Jose Maria Carazo, Andreas Plückthun, Ohad Medalia

## Abstract

The structural and functional characteristics of membrane proteins can be influenced by the composition of the membrane. Consequently, native membranes are most relevant for the study of receptors and other membrane proteins. In this study, we investigated two types of cell-derived vesicles: natively shed extracellular vesicles (EVs) and mechanically derived vesicles (MVs). To this end, we utilized the human breast cancer cell line SKBR3, which strongly overexpresses the receptor HER2. We designed a protocol based on designed ankyrin repeat proteins (DARPins) to purify EVs and MVs enriched in HER2, and to ensure the native orientation of the HER2 receptors within the vesicle. The isolated HER2-containing EVs and MVs were characterized by cryo-EM, cryo-electron tomography (cryo-ET) and mass spectrometry (MS), which revealed fundamental differences between the different vesicle types. Our study highlights the greater structural diversity of EVs over MVs. A single particle cryo-EM analysis and classification of all visible receptors on the vesicle surface yielded electron density consistent with HER2 at modest resolution. Taken together, our results suggest that MVs can serve better than EVs as a suitable platform for the structure determination of membrane proteins within their native membrane environments.

## Introduction

Membrane proteins are essential for cellular function, since they are involved in many fundamental processes, such as cell signaling, solute transport, immune responses, and synaptic transmission. Previous studies have shown that the local membrane environment intricately controls many aspects of membrane protein properties(*1, 2*). For example, the lipid composition of the membranes is coupled to protein conformational changes(*3, 4*), the membrane environment influences protein diffusion(*5*), and the interactions between co-localized proteins in the same membrane are linked with cellular signalling(*6*). Therefore, ideally, the structure of a membrane protein would be determined in its native membrane environment.

Structural determination of membrane proteins currently relies on overexpression systems(*7*) and the subsequent isolation of the proteins of interest in detergents, optionally followed by their incorporation into synthetic lipid bilayers(*8, 9*). This structural analysis provided unprecedented information expanding our understanding of how membrane proteins are functioning. However, the native membrane environment is lost during such sample preparations. Cryo-electron tomography (Cryo-ET) combined with focused ion beam (FIB) milling has emerged as a promising method for studying the structures of biological molecules in their native cellular environments (*10–13*). However, due to the dense molecular crowding in membranes, the small size of many membrane proteins and the low signal-to-noise ratio of cryo-ET, the structural features of many membrane proteins are challenging to interpret and/or too weak to be clearly identified and determined. As a result, this method is primarily limited to studying very large membrane-associated protein complexes, such as membrane attached ribosomes (*14*) or nuclear pore complexes(*15, 16*).

Cell-derived membrane vesicles have been recognized as promising platforms for studying membrane proteins in a close-to-native environment(*17–20*). Different types of membrane vesicles can be obtained, depending on the experimental procedure. For example, extracellular vesicles (EVs) are a heterogeneous group of membrane vesicles naturally released from cells(*21–23*). Different functions of EVs were considered, such as the selective elimination of proteins, lipids and nucleic acids from cells, but also intercellular communication via transfer of bioactive constituents from one cell to the other(*21, 24*). Another type of vesicles are mechanically induced membrane vesicles (MVs), also denoted cell-derived vesicles and a variety of other names(*25*). They have been prepared by extrusion through a syringe(*25–27*). They contain many of the original cellular lipids and membrane proteins, often in their original composition(*20, 28*). While different types of cell-derived membrane vesicles have already been used to investigate the structure of membrane proteins(*17, 29*), these studies were still built on protein over-expression procedures.

HER2 is a member of the EGFR family of receptors (*30*). Overexpression of HER2 has been implicated in the development and progression of tumors and is commonly associated with aggressive forms of breast and gastric cancers^6,7^. HER2 is the preferred dimerization partner of members from the EGFR family of receptors. As HER2 has no known ligand, it is always in the extended conformation able to dimerize with other EGFR members(*31*), and when highly overexpressed such as in several forms of cancers, it can hetro-dimerize even in the absence of a ligand of the other receptor(*32*).

HER2 has been recognized as a prime candidate for tumor-targeting therapeutics, since ∼20-30% of breast cancers and ∼20% of gastric cancers show HER2 overexpression, usually correlated with a rapid disease progression and poor survival rates^8-11^. Over the last few decades, small-molecule tyrosine kinase inhibitors (TKIs)^12-14^ and humanized monoclonal antibodies (mAbs)^8,15^ have been developed to target HER2-overexpressing tumor cells. While kinase inhibitors have resulted in low clinical efficacies during clinical trials^16-18^, probably because selection pressure leading to kinase mutations renders the inhibitor inactive, anti-HER2 mAbs have become an integral part of the standard of care for HER2-positive breast cancers. Indeed, combined therapies of mAbs with chemotherapy have turned out to be most efficacious in treating HER2-positive breast cancer^19-21^, and anti-HER2 antibody drug conjugates (ADCs) are being clinically used now as well(*33*).

Here, we set out to evaluate possible avenues to produce HER2-enriched vesicles. To this end, we utilized SKBR3 cells, a HER2-overexpressing human breast cancer cell line(*34*). Two types of cell-derived vesicles were produced and compared, namely spontaneously shed EVs and mechanically induced MVs. Using an affinity purification strategy based on designed ankyrin repeat proteins (DARPins)(*35*) immobilized on magnetic beads and cleavable by a specific protease, we successfully enriched HER2-containing vesicles. Cryo-ET and mass-spectrometry were employed to analyze the structure and composition of the vesicles, the crowdedness of their surface, and their membrane protein distributions. Finally, we applied cryo-EM on purified MVs to reconstruct receptor structures on the vesicle surface.

## Results

Naturally occurring extracellular vesicles are shed from the surface of cells into the extracellular space(*21–23*). They have drawn interest since they are altering the extracellular environment, participate in intercellular communication, and facilitate cell invasion through cell-independent matrix proteolysis(*36*). In human plasma, platelet derived EVs constitute the majority of the circulating vesicles found in blood(*37*). In some cancer cells, EVs are enriched in EGFR(*38*), and they were used to reconstruct a structure of EGFR dimers(*39*), although only to a modest resolution. On the other hand, mechanically induced vesicles (MVs) have been used to obtain insight into EGFR dynamics in cancer cells by NMR(*20*). More recently, mechanically induced vesicles were introduced for structural studies of over-expressed ion channels by cryo-EM(*29*). In the current work, we aimed to isolate and characterize the two different vesicle types, EVs and MVs for their suitability for studying the structure of natively expressed membrane receptors.

### Morphological characterization of EVs and MVs derived from SKBR3 cells

SKBR3-derived EVs were harvested and purified from spent media, obained from cells grown in serum-free medium for 14 h, while MVs were generated by passing cells through a syringe, a method shown to generate plasma membrane vesicles(*20*) (Fig. 1a and Methods section). To compare their suitability for structural analyses of receptors embedded in the plasma membrane, we purified EVs and MVs and subjected them to cryo-EM, cryo-ET, and mass spectrometry analysis (Fig. 1a). Low-magnification cryo-EM images of the EVs and MVs showed that the preparations exhibit similar characteristics and are both amenable for cryo-ET analysis (Fig. 1b,c). In this investigation, we acquired 35 and 36 tomograms from EVs and MVs, respectively. Subsequent inspection of the tomograms indicated variability and heterogeneity in the physical appearance of the vesicles. Examples of x-y slices through tomograms of EVs are shown in Fig. 1d. Based on their morphology, we classified 92 EVs into 9 structural groups: (i) Low inner-density vesicles (3.3%), (ii) elongated vesicles (3.3%), (iii) dense vesicles (8.8%), (iv) partially dense vesicles (22%), (v) cytoskeleton containing vesicles (13.2%), (vi) vesicles with droplet-like structures (droplets vesicles, 2.2%), (vii) multivesicular vesicles (multilamellar vesicles) (3.3%), (viii) crosslinked vesicles (4.4%), and (ix) others (39.5%) (Fig. 1e left).

**Figure 1.**
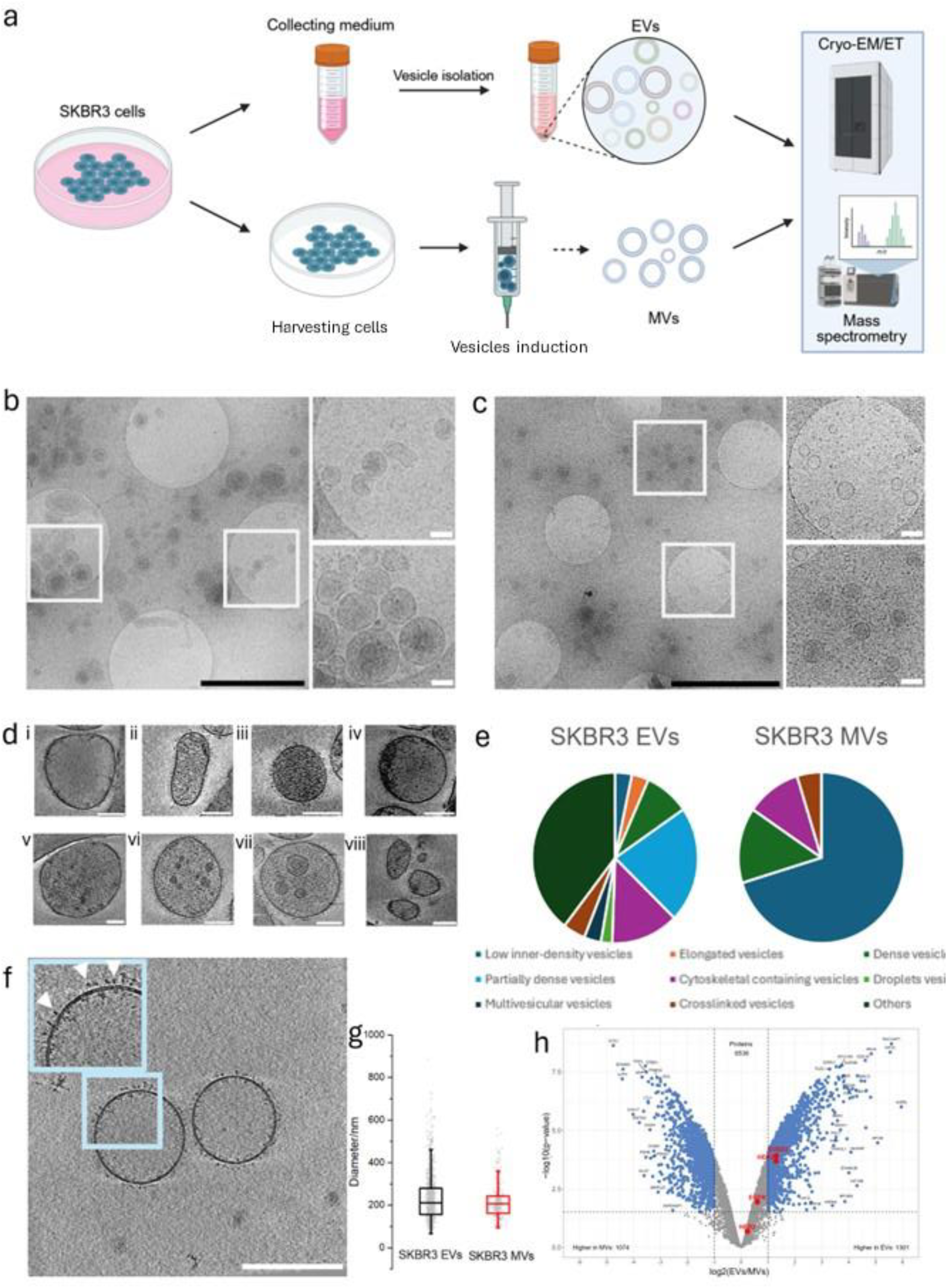
SKBR3-derived EVs and MVs. **a.** A schematic illustration of two different types of vesicle preparations and analysis. EVs are naturally released by cells, while MVs are mechanically created by extrusion through a syringe. **b** and **c.** Representative cryo-EM micrographs of SKBR3-derived EVs and MVs, respectively, indicating the comparable size of both vesicle types. White framed regions are shown at higher magnification (right). Black scale bars 2 µm. White scale bars 100 nm. **d.** SKBR3 EVs and MVs were classified into nine classes, based on their morphology (here exemplified by EVs): **i**-Low inner density vesicles, **ii**-Elongated vesicles, **iii**-Dense vesicles, **iv**-Partially dense vesicles, **v**-Cytoskeleton-containing vesicles, **vi**-Droplet-containing vesicles, **vii**-Multilamellar vesicles, **viii**-Crosslinked vesicles and ‘other vesicles’ (e.g., Fig. S8d). An *x-y* tomographic slice is shown for each class of vesicles. **e**. Pie diagrams showing the class distribution of EVs and MVs. The low inner density vesicles class is the most frequent class in MVs. **f.** An x-y tomographic slice through an SKBR3 MV. A higher magnification of the framed region is shown (top left), where receptors are visible (white arrowheads). **g.** The diameters of both EVs and MVs are comparable, although the MVs show less variability in size. **h**. Volcano-plot comparing the protein composition of SKBR3-derived EVs and MVs characterized by MS analysis. Most proteins showed non-significant differences in abundance between the two vesicle types. Members of the EGFR family (EGFR, Her2 and Her3) and SUSD2 is highlighted in red (More details in Supplementary Figure S2 b-f).

The MVs showed a comparable appearance to the EVs; in total, we analyzed 98 MVs and classified them using the same morphological classification criteria as for the EVs (Fig. 1e right). Cryo-ET analysis of MVs indicated electron densities emanating from the membrane, many of these likely represent receptors (Fig. 1f white arrowheads). Our classification indicated that 77.5% of the MVs appear round and lack substantial internal densities (Fig. 1e, right), resembling the low inner-density EVs, which suggests that MVs are morphologically more homogeneous than EVs. In addition, the diameters of MVs and EVs are comparable, although MVs show lower variance in their diameters (Fig. 1g)

To gain insight into the protein constituent in the spontaneously EVs and in the mechanically induced MVs, we performed a mass spectrometry-based proteome analysis of the two different vesicle types using a newly developed protocol for analysis of low sample loads(*40*). This analysis led to quantification of 6423 proteins from MVs and in quantification of 6607 proteins from EVs (averages of 3 bio-replicates). Among these proteins, 2375 proteins showed differential abundance between the two vesicle types (Fig. 1h and Fig S2). Looking closer into the proteins with different abundances in the two vesicle types it becomes clear that the shed vesicles fraction is richer in cytoskeletal proteins, Rab proteins and exosomal proteins, while the MV fraction seems richer in ribosomal proteins and proteins from the proteasome complex (Fig. S2). This could also be observed in cryo-EM images (see below).

Next, we applied the DeepLoc algorithm to predict the subcellular location of the identified proteins. The DeepLoc algorithm uses machine learning to predict the subcellular location of human proteins based on the protein’s amino acid sequence(*41*) . This analysis enabled us to predict which of the identified proteins that would locate into the cell plasma membrane, into organelles, the nucleus or if the proteins would be secreted. Focusing on the 10 % most abundant proteins indicated that the diversity of membrane proteins is higher in the EVs than in the MVs (Fig S2b). HER2 showed slightly higher abundance in EVs compared to MVs. In MVs, we detected similar levels of SUSD2 as of HER2.

We also used proteomaps (*42*) to visualize the proteome compositions of the two different vesicles preparations. It is noteworthy that MVs have a higher content of ribosomal proteins, proteasome-related proteins, and proteins involved in glycolysis compared to EVs (Fig S2h & i). The latter, however, showed higher abundance of cytoskeletal proteins and of proteins involved in signaling pathways. The observed proteome differences are attributable to the distinction between the pathway leading to EVs and the mechanical treatment of cells during the production of MVs(*21–23*),(*25–27*).

The comparison of the protein composition between EVs and MVs revealed a substantial discrepancy in their protein composition, despite the HER2 expression levels being similar in both SKBR3-derived EVs and MVs. Next, we proceeded to characterize the membrane protein composition, focusing exclusively on proteins containing signal peptides and transmembrane domains. This analysis demonstrated that HER2 is a highly abundant membrane protein in both of MVs and EVs. However, EVs exhibit slightly greater morphological and compositional heterogeneity compared to MVs, and have higher inner density, which is less favorable for cryo-EM based structural analysis.

### A general method for the purification of HER2-containing vesicles

During the production of MVs, the membrane curvature may become inverted, which results in an reversed orientation of the receptors, namely as “outside-in” instead of the native “outside-out” receptor orientation. Therefore, we developed an affinity enrichment strategy based on DARPins(*35*) immobilized on magnetic beads. Here, we took advantage of the DARPin G3, a previously developed HER2 binder that has a picomolar affinity to the extracellular domain (ECD) of HER2(*43, 44*), and could be equipped with a C-terminal protease cleavage site, followed by a C-terminal Avi-tag.

The process involved three major steps (Fig. 2a): (1) Streptavidin-coated magnetic beads were incubated with the biotin-conjugated DARPin G3 to produce G3-decorated magnetic beads; (2) binding of the HER2-containing vesicles to the DARPin G3-coated beads after mixing; (3) elution of the HER2-containing vesicles by proteolytic cleavage and release of the DARPin captured vesicles from the beads, followed by analysis by mass spectrometry and cryo-EM.

**Figure 2.**
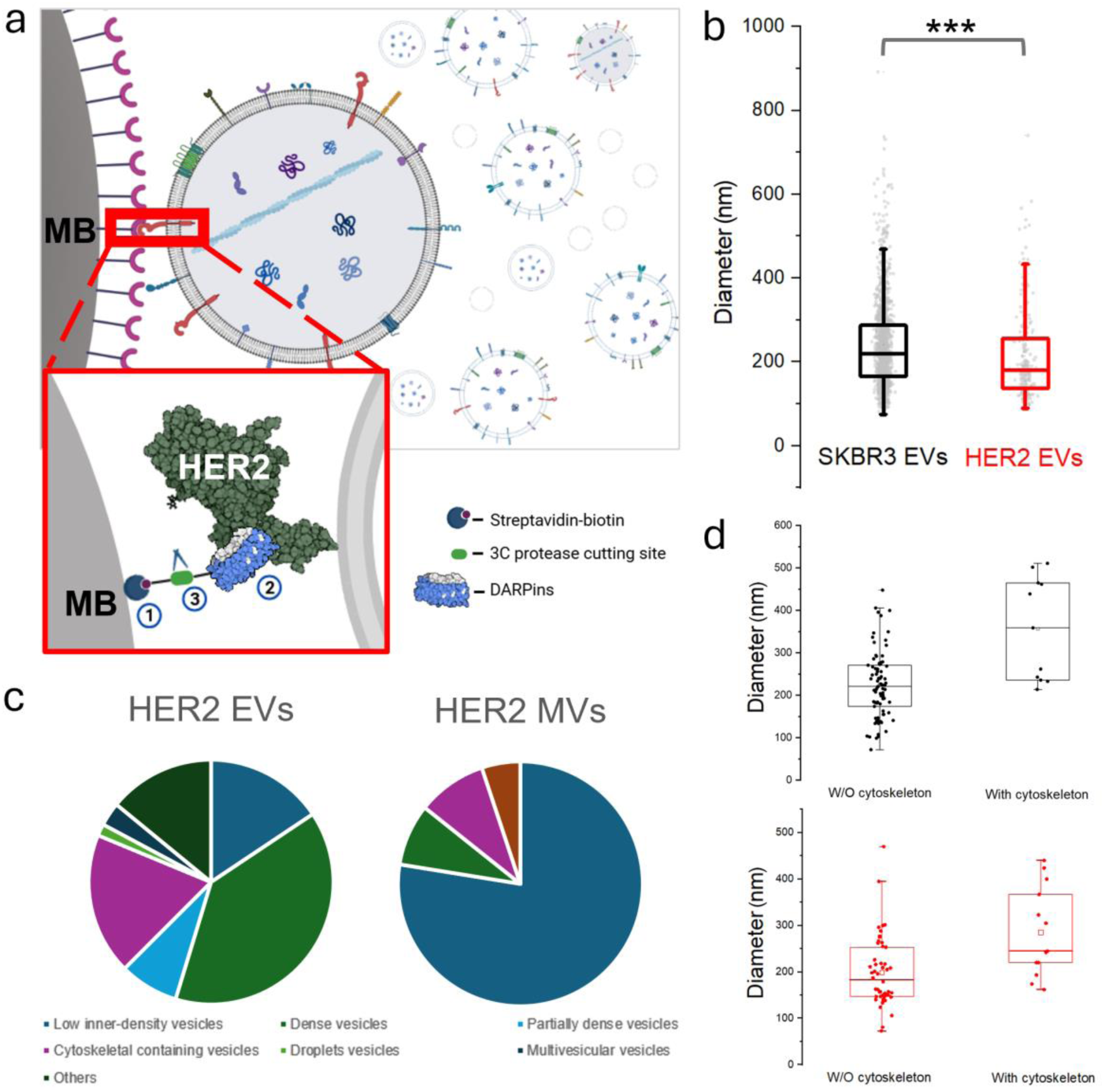
HER2-containing vesicles purified by magnetic beads coated with protease-cleavable DARPins. **a.** Schematic illustration of our approach towards affinity enrichment of HER2-containing vesicles. Biotinylated DARPins with affinity to HER2 were immobilized on streptavidin-coated magnetic beads (MB) ①. The biotinylation site is located on an Avi-tag, separated by a 3C protease cleavage site from the DARPin. ② Vesicle bound to the DARPin-coated bead via the HER2 ECD. ③ Using 3C protease, the immobilized HER2-containing vesicles were released. **b.** Enriched HER2-containing EVs showed a reduced diameter in comparison to non-enriched EVs. The diameter was 190 ±110 nm and 204±110 nm for HER2-enriched EVs and EVs, respectively (Average values are indicated by horizontal line, and the box range is from 25-75% of the whole population. **c.** The class distribution of EVs and MVs are displayed as a pie diagram. The enrichment approach influences the distribution of types of EVs more substantially than for MVs (cf. Fig. 1e). **d**. Size of EVs with (right) or without (left) cytoskeletal structures before (Upper panel) and after (Lower panel) HER2-based affinity purification, indicating the reduced size of the EVs independent of their content of cytoskeletal structures. MVs and HER2 MVs show no differences (data not shown).

The optimal conditions for binding biotinylated DARPin G3 to streptavidin coated magnetic beads and subsequent release by 3C protease cleavage were validated by SDS-PAGE analysis (Fig. S3). More than 50% of the G3 protein was biotinylated, as indicated by a shift to higher molecular weight (lane 2, band located at approximately 65 kDa) when forming an SDS-PAGE resistant complex with streptavidin. However, this shift was completely abolished upon treatment with 3C protease, which cleaves off the biotinylated Avi-tag (lane 4, Fig. S3). Since unbiotinylated G3 cannot bind to streptavidin beads, an additional washing after the first step ensures that unbound DARPins cannot interfere with the purification strategy. The feasibility of the approach was also validated using intact cells with or without HER2 expression (Fig. S4)

### Characterization of HER2-containing EVs and MVs

HER2-containing EVs and MVs were enriched using the strategy described above followed by cryo-ET analysis. Although the average diameter of HER2-containing EVs was found to be 190 ± 100 nm, and thus only slightly smaller than the average of the total population of EVs (Fig. 2b), indicated that this difference in diameter was significant. Next, we classified the enriched HER2-vesicles by their morphological characteristics, as done for the total EVs (Fig. 1d & e). Notably, two prominent vesicle classes observed in EVs (Fig. 1e) were absent from the HER2-enriched EVs (Fig. 2c): The classes of elongated vesicles and of crosslinked vesicles. In contrast, the class of dense vesicles increased in frequency from 8.8% in EVs to 39.1% among HER2-enriched EVs (Fig. 2c). Another notable change was that cytoskeleton-containing EVs appeared larger than those found in HER2-enriched EVs (Fig. 2d). A possible explanation is that the length of the cytoskeletal elements favors them to be present inside larger vesicles, which were underrepresented in the HER2-enriched fraction.

These substantial variations between HER2-enriched vesicles and the total EV population were not found in MVs, presumably since the major determinant of their size is their purification protocol and centrifugation step, and thus independent of the HER2 enrichment step. Therefore, the HER2-enriched MVs remained similar to the total population of SKBR3 MVs, with the low-density vesicles being the most abundant morphological class, accounting for over 70% of the vesicles in these samples (Fig. 2c).

Thus, significant differences were observed in the vesicle distribution both size-wise and morphologically between the naturally occurring EVs and the mechanically induced MVs. For instance, low inner-density vesicles accounted for only 15.6% of HER2-enriched EVs but for 70.2% of HER2- enriched MVs. Conversely, this resulted in a higher average inner density in HER2-enriched EVs compared to MVs (Fig. 2c), suggesting that EVs encapsulate a greater number of macromolecules. These differences imply that the release of EVs is not a random cellular process and that their structural characteristics may reflect specific biological functions(*45*).

A key advantage of the applied enrichment strategy is that both the native and the inverse orientation of HER2-containing vesicles can in principle be obtained, using DARPins that target the extracellular or intracellular receptor domains, respectively. However, in this study, we only used the DARPin G3 that binds to subdomain IV of HER2 ECD(*46*). The flexible attachment of this subdomain relative to subdomains I to III(*47*) allows the binding to the DARPin G3 even when this is attached to the large magnetic bead.

### Initial structure determination in native membranes

In many previous cases, structures of single trans-membrane-helix receptors were determined separately for the extracellular and the intracellular domains. However, this approach does not allow full visualization of the interaction with neighboring receptors in the same membrane or the effect of interacting proteins, e.g., therapeutic antibodies, or the impact of small molecules on the conformation of the receptor *in situ*. We therefore set out to develop strategies to allow structure determination of single trans-membrane receptors as full-length proteins in their natural environment. This is a challenging task due to several degrees of flexibility which characterize this group of receptors, even if dimerized(*48*). The HER2 receptor contains a flexible linker between the single TM-helix and its ecto-domain, allowing some free orientation relative to the plasma membrane(*49*). Furthermore, the different subdomains making up the extracellular domain can orient differently relative to each other. For example, even when purified, several conformations of the HER2 receptor have been detected, in addition to a spectrum of less populated conformations(*47*).

In the membrane of cell derived vesicles, many different membrane proteins can be expected, and this was indeed observed by our MS analysis (Supp. Table 1). This analysis showed that several proteins with similar molecular weight as HER2 are present in significant amounts and therefore may be misidentified as HER2 receptors, as the initial identification of protein complexes in cryo-EM and cryo-ET mostly depends on low-resolution information, defined by the size and the shape of the receptors. It is therefore extremely challenging to avoid that 2D and 3D classes are constructed by averaging more than a single type of membrane protein, which would be detrimental for the structure determination of the targeted receptor residing in the membrane.

As expected, cryo-ET analysis of HER2-containing EVs and MVs revealed a high number of proteins with similar dimensions as HER2 (Fig. 3a & b). In these tomograms a better contrast for MVs than EVs was observed as a result of the lower intra-vesicular density of the MVs (Fig. 3b & c). Although SKBR3 cells are known to natively express a very high copy number of HER2 receptors, the densely packed plasma membrane of the cells highlights the complexity of native membranes and the diversity of proteins embedded in the membranes. Thus, the high density of proteins on the surface of MVs indicated another challenge which comes from the fact that HER2 receptors are localized in proximity of many other proteins, and they may even interact with additional membrane proteins in their native environment.

**Figure 3.**
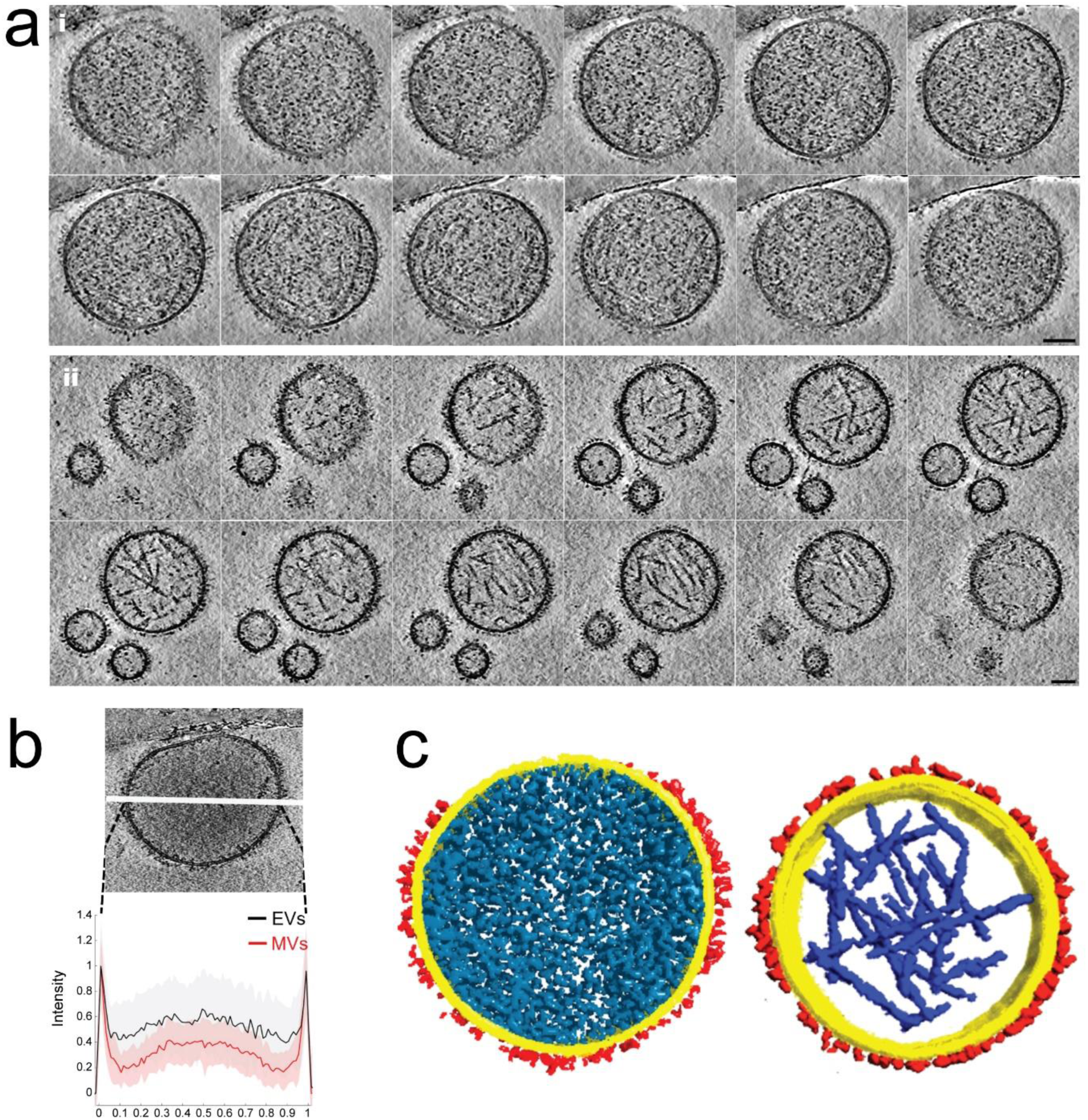
HER2-containing EVs and MVs. **a.** x-y tomographic slices through representative HER2-containing EV (i) and MVs (ii). Scale bars is 100 nm. **b.** The average line density through HER2-containing EVs and MVs indicated that the MVs are less dense. The standard deviation is shown by the shadowed area. The central line is drawn (top image, with a slice through an EV in the top half of the picture and a slice through a MV in the bottom half). **c.** The HER2-enriched EV (left) and MV (right) are shown as rendered views. Membrane (yellow), receptors (red), cytoskeleton (blue) and molecular structures (cyan).

While sub-tomogram averaging(*50*) may potentially be used to determine structures in the isolated vesicles, this method so far showed only limited success in resolving structures of receptors embedded in plasma membranes(*51*). In contrast, high resolution structures of ion channels have been resolved from cell-derived membrane vesicles(*29*). Importantly, the resolved structures were very homogeneous, probably due to the substantial rigidity of the membrane-embedded part of these proteins. Furthermore, they exhibit a molecular weight more than twice that of the HER2 receptor (138 kDa). Nonetheless, these results suggest that cryo-EM could potentially be the method of choice for determining the structure of receptors embedded in MVs. To test this hypothesis, more than 100,000 micrographs were collected from SKBR3-derived MV samples. The high image quality of these rather thick specimens was validated by determining the structure of ribosomes found in the samples. Indeed, a well-resolved ribosome structure was obtained (Fig. S5, 2.8 Å resolution), confirming the quality of the dataset and its high-resolution structural information, which is present in the data.

To address the expected diversity among proteins included in our data from the analyzed particles, we first analyzed what the mass spectrometry results indicated as the most abundant membrane proteins in MVs using AlphaFold(*52*) (Fig. 4a). We found that HER2 was the only abundant membrane protein equipped with both extracellular and intracellular domains that would be large enough to be detected in cryo-EM micrographs. Furthermore, based on a previous structure of HER2, we expected that the HER2 extra-cellular domain (ECD) would extend approximately 10 nm away from the membrane, while the HER2 intra-cellular domain (ICD) would extend approximately 3 nm away from the membrane (PDB:1N8Z, 3PP0)(*53*). However, the conformational heterogeneity of HER2 may impact these distances, due to the inherent flexibility of full-length HER2 and especially the hinges connecting the transmembrane helix on either side to the domains outside the membrane(*49*).

**Figure 4.**
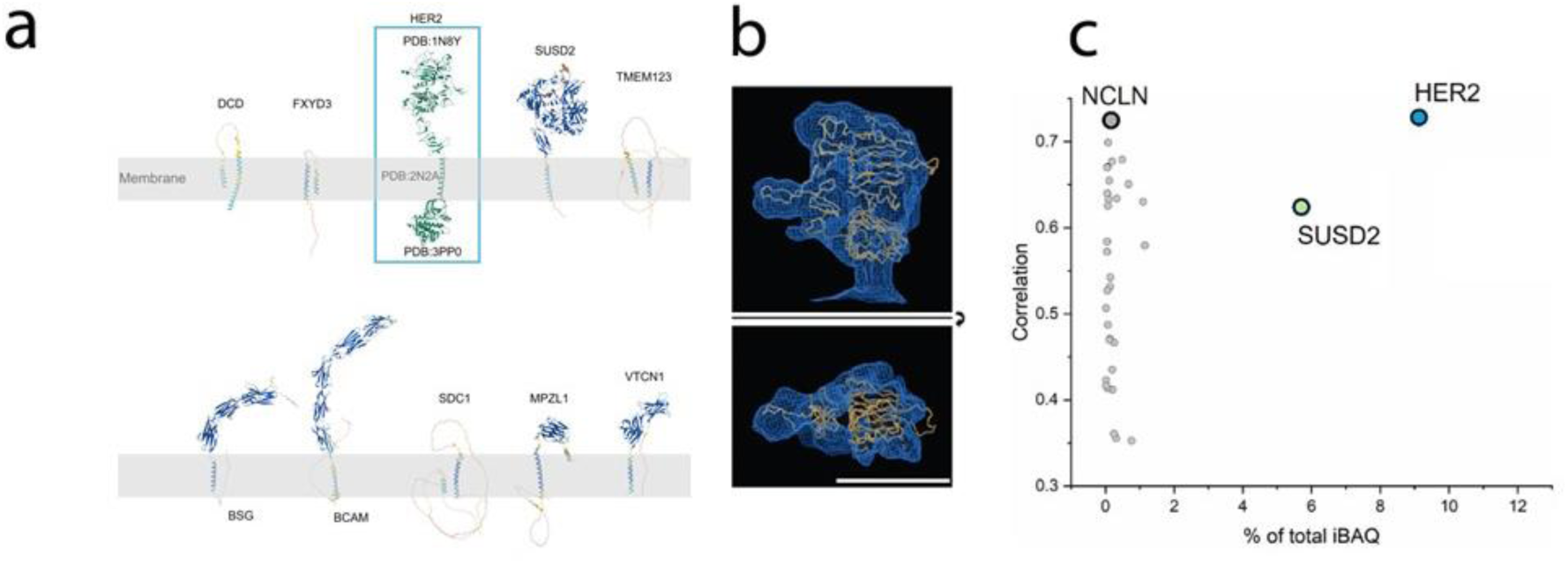
Structure determination of receptors from SKBR3-derived MVs. **a.** The most abundant membrane proteins (by mass spectrometry analysis) in MVs were modelled using AlphaFold2, indicating that most of the abundant proteins are structurally distinguishable from HER2. **b.** The derived HER2 structure docked into the calculated 3D structure, fits well. The threshold of the structure was increased to also include less rigid features. The structural subdomains of I, II and III of HER2 ECD were used for the docking. **c.** Visualization of the structural correlations between 34 membrane proteins found in the vesicles, which have similarly sized non-membrane domain as HER2, with the resulting 3D density map for HER2 are shown as a function of the estimated protein abundance (total iBAQ). While HER2 showed the highest correlation, the ER protein NCLN showed a similar correlation value, however this membrane protein is found only in limited amounts. The protein SUSD2 appeared in a similar—lower though—abundance compared to HER2, but it showed only a correlation of 0.62, much lower than the reconstructed structure.

Therefore, we chose an image processing workflow that is guided by prior structural knowledge, i.e., the domain organization of HER2. After identification of all vesicles in our data set, using the distinctly detectable lipid bilayer as a template (Fig. S6a), all electron densities emanating from the membrane were picked using a manually labeled Topaz(*54*) model. False positives, primarily located at carbon edges or inside vesicles, were removed based on the absence of bilayer structural features (Fig. S6b,c). Additional filtering of falsely picked densities was achieved by applying the expected dimensions of the HER2 ECD and ICD as criteria. Next, initial 2D classification was performed to exclude densities smaller than 5 nm in diameter, which are unlikely to correspond to the HER2 ECD (Fig. S6d,e).

This procedure resulted in 151,480 particles with substantial densities emanating from the membrane. Next, we analyzed the ICDs that are located in proximity to the ECD, although their position may not be fixed, given the expected flexibility of the cytoplasmic region of the HER2 receptor, both regarding rotation and position. To classify the ICD independently, the coordinates of each particle were shifted to the inner side of the membrane, and the particles were re-extracted, followed by several 2D classification steps. All classes that displayed potential ICD densities were selected, which resulted in 55,760 particles (Fig. S6f,g). These particles were used to reconstruct a 3D map to modest resolution, which was subsequently used as template for a 3D classification performed in RELION(*55*), selecting ∼50% of the particles as HER2 receptor candidates (Fig. S6h). This resulted in a structure resolved to 8.7 Å (Fig. 4b).

To analyze this structure, HER2 domain I-III was fitted into the cryo-EM map using the Phenix package(*56*) (Fig. 4b). Although HER2 has not previously been determined within its native membrane, it is well accepted that subdomains I-III of its ECD are structurally rigid(*47*). Therefore, a model consisting of HER2 subdomains I-III was used. The reconstructed structure clearly resembles the shape and size of the HER2 ECD resolved to <9 Å (Fig. S7). However, the refined structure did not show a clear ICD structure (Fig. S7). Similarly, in the structural analysis of purified quasi-full-length HER2, the flexibility in orientation of the ICD also prevented its structure determination, except at very modest resolution(*47*). Interestingly, several structural classes obtained in the present analysis of sub-tomograms extracted from the vesicles resemble a dimeric assembly of EGFR family of receptors (Fig. S8). However, the limited number of detected structures in this conformation prevented the 3D reconstruction of these potential dimeric receptors.

To provide additional insight into the interpretation of the densities observed at the SKBR3 membrane surface, we analyzed structures predicted with AlphaFold2(*52*) for all high abundance membrane proteins detected by mass spectroscopy and compared them with the reconstructed maps. We then systematically compared these predicted structures with the final density maps, after categorizing the membrane proteins into six groups (Fig. S8c and Table 1). We eliminated 5 of the 6 groups as candidates for representing HER2: (i) low-abundance proteins that represent <0.01% of the total membrane protein content. This group of proteins was excluded from further analysis, because the low copy number (estimated as Intensity-Based Absolute Quantification, iBAQ, values from the MS data) reduces the probability that these proteins represent a large enough population to be reconstructed; (ii) proteins with large extra-cellular membrane domains. This group of membrane proteins contains ECD domains larger than 1000 amino acids, which were excluded due to the incompatibility with the reconstructed map; (iii) proteins with small ECD domains (<450 amino acids) were also excluded from further analysis. (iv) proteins with tandem repeat domains. This group of proteins includes 26 membrane proteins, most of which are involved in cell adhesion or intercellular junctions. All of them have a unique structural feature which differs clearly from our map. Moreover, similar structures were observed in tomograms, where vesicles are crosslinked with each other (Fig. S8d); (v) disordered proteins: AlphaFold2(*52*) predictions for these 13 membrane proteins were too disordered to allow meaningful comparisons.

After eliminating the proteins from the five groups, the above procedure resulted in 34 remaining membrane proteins that could be misinterpreted as HER2 (Supp. Table 1, bottom). Therefore, we downloaded their full length AlphaFold2(*52*) structures and trimmed the flexible domains based on the pLDDT confidence score. We then calculated the correlation coefficients (CC) for each structure with our HER2 density map, using UCSF Chimera(*57*). Here, subdomains I-III of HER2 ECD showed the highest CC value (CC = 0.724) with the derived structure followed by the ER protein NCLN (CC=0.720) present only in very low copy numbers (by iBAQ value estimation). In contrast, a plasma membrane protein with similar--lower though--abundance compared to HER2 showed a significantly lower CC value (CC = 0.623). These results support the workflow, but cannot exclude that some included electron densities may be “contaminated” with electron densities from other protein species, such as NCLN and SUSD2. It is likely, however, that the majority of the density was contributed by the highly expressed HER2, and that it is the HER2 receptor that was reconstructed from the native plasma membrane of cell-derived MVs, although at modest resolution.

## Discussion

In the work presented, we assessed the potential of using different vesicle types derived from SKBR3 cells for the structural determination of HER2, which is highly expressed on SKBR3 cells. The naturally shed EVs offer a promising experimental system for studying receptors in vesicles, particularly due to the relatively high levels of HER2 receptors present in these vesicles. However, the EVs released by SKBR3 cells appeared to be heterogeneous in size and exhibited a relatively dense appearance under cryo-EM analysis. This characteristic rendered the sample less electron transparent, which is a prerequisite for high-resolution cryo-EM and tomography analysis. Notwithstanding, given the elevated HER2 expression levels, this system may offer significant advantages for biochemical analysis, such as measurements of binding affinity of protein ligands, but may be less well-suited for structural investigations.

In contrast, our findings showed that mechanically induced vesicles (MVs), exhibit greater homogeneity in terms of morphology and size distribution, compared to EVs, while the expression levels of HER2 were found to be similar. Cryo-ET analysis highlighted the distinctive features of these vesicle preparations. However, the sub-tomography averaging procedure did not yield a satisfactory reconstructed 3D structure. This was presumably due to the relatively low molecular weight of the HER2 receptor (<200 kDa), its orientational flexibility, the high contrast of the membrane, and the low signal-to-noise ratio of the tomograms. The application of cryo-ET to individual proteins, such as HER2, would facilitate the attainment of a more reliable 3D classification, allowing the identification of the complexes under scrutiny. Consequently, the image processing workflow employed in this study focuses on MVs using single particle analysis (SPA) for structural analysis.

Recently, several approaches have been developed for resolving heterogeneity in SPA, which demonstrated efficacy for purified protein complexes(*58, 59*). Previous studies have also examined the application of cryo-EM for analysis of purified vesicles with over-expressed membrane proteins, focusing on protein with a rather rigid transmembrane region. Nonetheless, several challenges become evident when the aforementioned methods are applied to receptors with single transmembrane helices, which ultimately compromise the precision of the analysis when vesicles derived from native membranes are utilized. The primary challenge is the presence of a substantial number of other membrane proteins in native MVs in addition to the protein of interest, as evidenced by MS analysis. This results in a diverse array of membrane proteins that are picked for the classification and subsequent analysis. Consequently, the selected particles likely included other membrane proteins with similar low-resolution structural features. This could not be overcome despite the utilization of prior knowledge, such as predicted structural models of the protein of interest and of the other highly abundant membrane proteins present in the sample. Nonetheless, the approach facilitated a further computational enrichment of HER2 receptors in the particle picking step. In this regard, the development of cryo-EM labeling systems(*60, 61*) may facilitate circumventing this limitation in subsequent studies, which are under way.

An inherent challenge in the structural studies of single-transmembrane helix receptors like HER2 is the variability in ECD orientations. This flexibility of the angle between subdomain IV and the transmembrane helix has also been demonstrated previously for other EGFR-related proteins(*47, 48, 62*). A challenge with both vesicle preparations is the relative large size of the vesicles as cryo-EM specimens. The mean diameter of the vesicles exceeds 150 nm, thereby leading to augmented ice thickness. In comparison, high-resolution structures of the ion channel Slo1 were resolved from vesicles with a diameter of approximately 40 nm (*29*), a size obtained by sonication. Thicker ice will result in reduced signal-to-noise ratio in acquired micrographs, which in turn, will have a direct impact on the resolution of determined structures. Despite thicker ice, the samples permitted determination of ribosome structures at high resolution (Fig. S5), the reconstruction of small and more fragile elements, though, remains challenging.

Although the resolution of our reconstructed structure was insufficient to produce a high-resolution density map of HER2, a thorough comparison with other proteins in our sample strongly suggested that the structure which was reconstructed is the receptor. Although a high-resolution structure of HER2 could not be derived, the approach followed suggests that a human breast cancer cell line like SKBR3, can be used for structural analysis. The ICD part of HER2 becomes much weaker after reconstruction, which indicate high flexibility between the ECD and ICD of HER2 with native membranes, as found in previous studies (*47, 63*). Nonetheless, a dedicated averaging of micrographs focusing on this domain may help to gain further information in these structural features.

In summary, the present study reports the differences between EVs and MVs for structural studies, thereby evaluating the applicability of these preparations as specimens for *in situ* membrane protein structural analysis. Future advancements in cryo-EM and tomography are anticipated to improve high-resolution structural analysis of HER2 in its native environment and so will also further developments in labeling technology. Such technological advances hold the potential to enable the reconstruction of individual transmembrane receptors within cellular contexts.

## Acknowledgements

Imaging was performed with equipment maintained by the Center for Microscopy and Image Analysis (ZMB), University of Zürich. We thank R. Boujemaa-Paterski for helping with the artwork. This work was supported by the European Research Council grant HighResCells (ERC-2018-SyG, proposal: 810057 to J.O, J.M.C. A.P., and O.M.).

## Author contributions

Cloning, expression, purification, cryo-EM sample preparation, and cryo-EM data acquisition were performed by C.W. with the help of G.N.D and M.E. Receptor reconstruction analyzed and interpreted cryo-EM data by C.W. under the supervision of M.E. MS data acquisition and analysis was conducted by O. Ø. Project management was carried out by J.O., J.M.C., O.M., and A.P. The initial manuscript was prepared by C.W., O.M. and M.E. All authors contributed to the result interpretation and discussion, as well as to the final editing and approval of the manuscript.

## Methods

### Cell culture

SKBR3 cells and CHO cells were cultured in Roswell Park Memorial Institute (RMPI) 1640 medium (GibcoTM, 21875-034) containing 10% (v/v) Foetal Calf Serum (FCS) (BioConcept, 2-01F10-I) and 1% (v/v) penicillin-streptomycin (Sigma-Aldrich, P0781) at 37 °C and 5% CO_2_. For fluorescence analysis, the cultured cells were first detached with 1x trypsin-EDTA solution (Sigma-Aldrich, T4174) and then seeded onto a 35-mm cell culture dish with glass bottom (MatTek, P35G-1.5-14-C) and grown overnight.

### EVs and MVs vesicle purification

To prepare extracellular vesicles (EVs), 10 × 15 cm cell dishes of SKBR3 cells at approximately 80% confluence grew in serum-free RMPI 1640 medium overnight. The spent medium was collected (∼200 mL), and cells and debris were removed by centrifugation at 2000 × g for 10 min. The supernatants were then filtered using 0.8 μm filters (Sartorius Stedium Biotech GmbH) before concentration with Amicon Ultra - 15 centrifugal filters (10 kDa) to reduce the volume to ∼10 mL. EVs were pelleted by centrifugation at 4°C for 30 min at 20,000 × *g*. The supernatants were carefully removed and 1 mL of PBS was added for resuspending the EVs, which were then pelleted again. The enriched EVs were suspended in 20 µL PBS for characterization.

To prepare mechanically induced vesicles (MVs), 10 × 15 cm cell dishes of SKBR3 cells at approximately 80% confluence were used. The cells were manually detached with a cell scraper and suspended with homogenization buffer (20 mM Tris pH 7.4, 1 mM EDTA and 250 mM sucrose, supplemented with freshly added phosphatase inhibitors and a protease inhibitor cocktail (Complete Mini, Roche). Syringes (27G × 1½, 0.4 mm × 40 mm) were used for generating vesicles from the cells. Subsequently, the unbroken cells and cell debris were removed by centrifugation at 1,000 × g for 10 min followed by centrifugation of the supernatant at 10,000 × g for 50 min at 4°C. The membrane vesicles were then collected at 110,000 × g for 1 h at 4°C and resuspended in 20 mM HEPES (pH 7.4) for characterization.

### Specific binding of magnetic beads to HER2 expressing cells

SKBR3 cells and CHO cells—not expressing HER2—were cultured in 12-well cell culture plates. Aliquots of 10 µL magnetic beads were coated (see next section) with DARPin G3 or E3-5 (with no measurable affinity for HER2 as a control) prior to incubation with cells in 1 mL serum-free RPMI 1640 medium for 1 h. To avoid bead sedimentation, the cell plates were gently shake at 100 rpm/min. Then the cells were washed 3 times for 10 min with a higher shaking speed of 200 rpm/min. Light microscopy images were acquired to verify the specificity of the binding.

### Enrichment of HER2-positive vesicles using magnetic beads

To enrich for vesicles that expose the extracellular domain of HER2, HER2 binding DARPins with a 3C cleavage site (described below) were coupled to magnetic beads for vesicle enrichment. The 3C cleavage site enabled easy release of the vesicles by addition of the 3C protease.

The DARPin G3 (abbreviated as G3)(*43*) was fused with an N-terminal MRGS-His_6_ tag and a C-terminal Avi-tag preceed by a 3C protease cleavage site (LEVLFQGP). G3 was expressed and purified as described previously(*43*). The DARPins were biotinylated in vitro using biotin ligase and purified using Nickel-NTA and gel filtration chromatography. The protein concentration of purified biotinylated DARPin G3 was 294 μM, evaluated with a Nanodrop 2000 Spectrophotometer (Thermo Scientific) at 280 nm.

The biotinylated DARPin G3 was then immobilized on streptavidin-coated magnetic beads. For each experiment, 10 μL of G3 were incubated with 200 μL Dynabeads^TM^ Biotin Binder (Invitrogen^TM^, 11047) for 1 h, at 4°C. After 3 washes (10 min) with 20 mM HEPES 1% BSA, the G3-coated beads were incubated with SKBR3 cell-derived vesicles for 1 h at 4°C. Then the beads were washed three times with 20 mM HEPES buffer to remove unbound vesicles and pelleted by a homemade weak magnet to minimize bead aggregation. After the first two washes, the beads were resuspended in 20 mM HEPES with 1% BSA, and after a third washing step, the beads were resuspended in 20 mM HEPES without BSA. Vesicles were then released from the beads by addition of 4 μL 3C protease (GenScript, Z03092) followed by incubation for 2 h at 4°C. The supernatant was collected and concentrated using a Vivaspin-500 filter (100 kDa, Sartorius, VS0141) to ∼10 μL for cryo-EM analysis. For the control experiment, a HER2-non-binding DARPin (E3-5) was used at 241 μM, and the 11 μL of E3-5 in the same format was used for immobilization on magnetic beads in each individual experiment.

### Immunofluorescence and confocal microscopy

Cells cultured on coverslips were fixed using 3.7% paraformaldehyde (4 min) and then permeabilized with 0.1% Triton X-100 (Sigma-Aldrich, T8787) for 10 min prior to incubation with blocking buffer (2% w/v BSA, 22.5 mg/mL glycine in PBS, pH 7.5, supplemented with 0.1% Tween-20 (PBST)) for 1 h at room temperature. The primary anti-HER2 antibody 3B5 (Merck, MABE330, 1:1000) was diluted in blocking buffer and incubated with the cells overnight at 4°C. After 3 washes with PBST, the FITC-labeled secondary antibody (Jackson ImmunoResearch, 715-095-150, 1:200) was applied for 1 h at room temperature. Cell nuclei were stained with Hoechst 33342 (Sigma-Aldrich, B2261) at room temperature for 30 min. The samples were then further washed (3x10 min) with PBS. HER2 localization was recorded in the GFP channel, and the Hoechst channel was recorded to visualize nuclei. The fluorescence images were acquired using a DMI4000B microscope (Leica Microsystems), and confocal scanning microscopy images were acquired with a Leica SP8 inverse FALCON laser scanning microscope. 3D stacks were reconstructed in Imaris 9.6.0.

### Mass spectrometry analysis of vesicle samples

#### NTA measurements

Vesicle samples with EVs and MVs were analyzed by nanoparticle tracking analysis (NTA) using a Zeta View PMX-220 NTA analyzer (Particle Metrix , Germany) equipped with a 520 nm laser to estimate the particle concentrations. In brief, samples were diluted with PBS to a of 10,000 fold final dilution. From this dilution, 1 mL was injected into the NTA analyzer and nanoparticles were counted by scanning at 11 different locations twice at 22 °C. The recorded data were analyzed and summarized using the ZetaView software version 8.05.14 SP7 reporting vesicle concentrations in the original samples.

#### OneTip sample digestion and desalting

For tryptic protein digestion prior to analysis by mass spectrometry, EV and MV samples were digested directly on EvoTips, essentially as described (*40*). In brief, samples were diluted to 10^7^ particles/µL. Five µL diluted sample (equivalent to 50 million particles) were then loaded on EvoTips already prepared as follows: First, tips were rinsed with 20 µL 80% acetonitrile, 0.1% TFA, then soaked with isopropanol, followed by a step to equilibrate the C18 material by loading 20 µL 0.1 % formic acid. After each step the reagent was removed by gentle centrifugation of the tips in the EvoTip box at 600xg for 60s. The equilibrated tips were then supplied with 5 uL digestion mix (100 mM triethyl-ammonium bicarbonate, 0.2% n-dodecyl-β-D-maltoside (DDM), 20 ng/μL Trypsin, and 10 ng/μL endo-LysC) before addition of 5 uL sample with EVs or MVs. Mixing was ensured by centrifugation for a few seconds at 50xg before incubation at 37 °C in an EvoTip box filled with just enough water to ensure that the C18 part of the EvoTips were immersed into the water. After 4 hrs, 50 µL 0.1 % formic acid were added to the EvoTips followed by centrifugation 60s at 600xg. Then the tips were desalted by washing with 20 µL 0.1 % formic acid before adding 100 µL 0.1 % formic acid and placement in an EvoTip box for running the sample using an EvoSep One system.

#### Mass spectrometry analysis

Samples were eluted directly from EvoTips onto a C18 column (Aurora Rapid75, 5 cm x 75 µm, 1.7 µm beads, IonOptics) and eluted into an Astral mass spectrometer (Thermo Fisher Scientific) using an Evosep One LC-system (Evosep, Odense, Denmark) running 16 min gradients (80 samples per day). The spray voltage was 1800V and the ion transfer tube was 275 °C. Full scan spectra (380-980 Th) were recorded at 240,000 resolution, 500 % AGC, max 30 ms injection time, while MS2 spectra were recorded in a data independent acquisition mode by 149 sequential narrow windows (4 Th), max 6 ms injection time, a normalized collision energy at 25 and an RF lens set to 40%.

#### Database searching and protein quantitation

The generated raw files were searched using Spectronaut v20.2 against the SwissProt database including 20,426 human sequences downloaded on 17-10-2023) along with small FASTA files containing sequences for DARPin 9.26, streptavidin, 3C protease, trypsin and endo-lysyl endopeptidase (endo-LysC) and common contaminants from bovine serum often observed in samples from cell culture. The following settings in Spectronaut were applied: Enzyme: Trypsin/P, peptide length - min 7 AA, max 52 AA, max 2 missed cleavages; Fixed modifications - none; Variable modifications - Oxidation(M), Acetyl (Protein N-term); Protein LFQ method: MaxLFQ, Quantity level: MS2, Quantity type: Area. iBAQ values were calculated outside Spectronaut by dividing MaxLFQ quantities from Spectronaut by the *in-silico* calculated number of observable tryptic peptides with no missed cleavage sites.

### Cryo-ET and cryo-EM, sample preparation and data acquisition

Tilt series were acquired using glow-discharged holey carbon copper EM grids (R2/1, 200 mesh, Quantifoil). 3 μL vesicle samples were introduced to the EM grids, prior to addition of 3 μL of 6 nm gold fiducial markers (Aurion). Samples were freeze plunged using a Vitrobot (Thermo Fisher) for snap freezing. Tomograms were mainly acquired using a FEI Titan Krios G1 electron microscope (Thermo Fisher) equipped with a Gatan Quantum Energy Filter and a K2 Summit direct electron detector (Ametek). Data were acquired at 81k magnification, resulting in a pixel size of 0.175 nm. All tomograms were acquired at 2.5-3.5 μm underfocus, covering an angular range of ±60° with a 3° increment, and a total electron dose of ∼120 e^-^/Å^2^. SerialEM was used for tomogram acquisition and IMOD(*64*) was used for reconstructions of tomograms.

A total number of 37, 56, 144, and 103 tomograms were collected from the samples with EVs, HER2-enriched EVs, mechanically induced vesicles (MVs) and HER2-enriched MVs, respectively. Diameters of the vesicles were manually measured in 3dmod and analyzed in Microsoft Excel and Origin (OriginLab Corp.).

For single particle cryo-EM, 3 μL vesicle samples were loaded on glow-discharged holey carbon gold EM grids (R2/2, 200 mesh, Quantifoil). The grids were blotted briefly, and an additional 3 μL vesicle samples were loaded, in order to increase the vesicle densities on the grids. All EM grids were vitrified using a manual plunger (MPI, Martinsried). The data were collected using a Titan Krios G3i (Thermo Fisher) equipped with a Gatan BioQuantum Energy Filter and a K3 Summit direct electron detector (Ametek), at a magnification of 130k, resulting in a pixel size of 0.651 Å. The micrographs were automatically acquired with an underfocus range between 1.5 -3.2 µm using Thermo Fisher Smart EPU software with a total electron dose of ∼60e^-^/Å^2^.

### Quality control of the data by ribosome reconstruction

Ribosomes present in the sample, enclosed by vesicle membranes, were used to confirm the quality of the samples. The image processing of ribosomes was performed in Scipion(*65*). A dataset of 12,373 movies of MVs was used. The particle picking was divided into two steps: the first step was the manual picking of ribosomes in a few micrographs, and the second step the auto-picking using the xmipp software. The CTF was estimated using Gctf. A total of 257,176 particles were extracted at 1.3 Å/pixel in a box-size of 400×400 pixel^2^ and used for 2D classification using CryoSPARC(*66*). 87,891 particles were used for the initial model, generating a 3D map using CryoSPARC. After using non-uniform refinement in CryoSPARC, a map at 2.8 Å resolution and B-factor -45.3, was obtained. The sharpening was applied to the final map, indicating that the sample allows to obtain structures at ∼3 Å.

### Image processing of MVs

All vesicle micrographs were imported into CryoSPARC(*66*). In the software package, the Patch Motion Correction, Patch CTF Estimation, Blob Picker and Topaz Extract were used for frame alignment, CTF correction, and particle picking, respectively. For the primary classification to remove false particles located on carbon edges or inside vesicles, the particles were extracted in a box size of 48 × 48 pixels with binning 10. For the second classification with the aim to detect low-resolution structural features of membrane proteins, the particles were extracted in the same box size but with binning 6. The ab-initio reconstruction procedure was selected for initial template creation and used for primary 3D particle alignment with non-uniform refinement. The particle center was shifted to the inside of the vesicles with the volume alignment tools, and for the intracellular part, the particles were re-extracted in a box size of 34 × 34 pixels with a binning factor of 6. After 2D classification, the whole map was reconstructed with the particles extracted in a box size of 84 × 84 pixels with binning 6, while maintaining the primary alignment.

To transfer the particles from CryoSPARC(*66*) to Relion5.0(*55*), the particles were extracted in a box size of 80 × 80 pixels with binning 3 in CryoSPARC(*66*), and then the PYTHON script csparc2star.py(*67*) was used for the conversion to the RELION format.

The 3D classification and 3D refinement were performed in Relion5.0(*55*). The template was derived from the previous non-uniform refinement map, with C20 symmetry applied. Blush regularization(*68*) was used for 3D classification.

### Molecular docking of AlphaFold models into the experimental density map

The AlphaFold2(*52*) predicted structures of full length HER2 were downloaded. The truncated structure consisting of subdomains I, II, and III was fit to the density map using Phenix docking(*56*). The AlphaFold2(*52*) predicted structures of all other abundant and possible membrane protein candidates were also downloaded. According to the domain boundaries and the model confidence derived from the pLDDT score, only the domains which are located outside of the membrane were used for docking. The models were fitted into our experimental 3D data and the correlation coefficient was recorded in Chimera(*57*).

### Statistical analysis

Segmentation of vesicle tomograms was conducted in the Dragonfly 3D software. Analysis of internal density of vesicles was calculated in MATLAB using the TOM Toolbox(*69*).

## Data availability

The recorded mass spectrometry RAW files along with the search result has been uploaded to the data repository PRIDE (*70*)and are accessible with the following identifier: PXD074163

The output from Spectronaut were analyzed further using Rstudio (v2022.12 B353, R version 4.2.2). Proteomaps(*42*) were used to visualize the quantitative composition of selected samples.

