## Supplemental Figures for "Comparison of extracellular vesicles and mechanically induced vesicles for structure determination of membrane proteins"

### Supplementary figures and tables

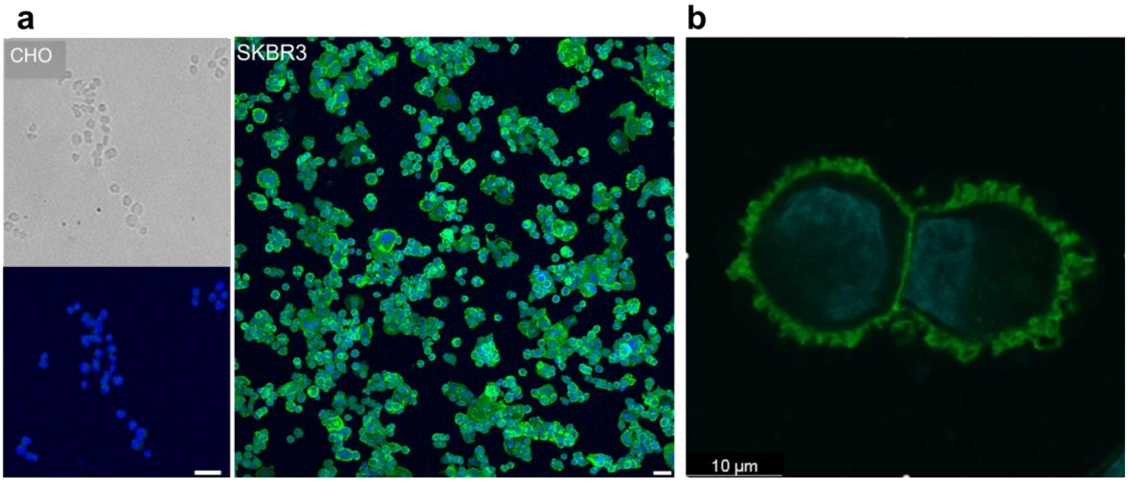


**Figure S1. Immunofluorescence microscopy images showing presence of HER2 in the human breast cancer cell line SKBR3. a.** A strong HER2 signal (Green) is detected in SKBR3 cells, using anti-HER2 antibody (Merck, MABE330), but no signal was detected in the control CHO cells (bright field and fluorescent channel are shown, up and bottom, respectively). Scale bar 40μm. **b.** The cellular ruffled plasma membrane of SKBR3 cells is revealed by confocal microscopy imaging. Chromatin (blue) stained using bisbenzimide.

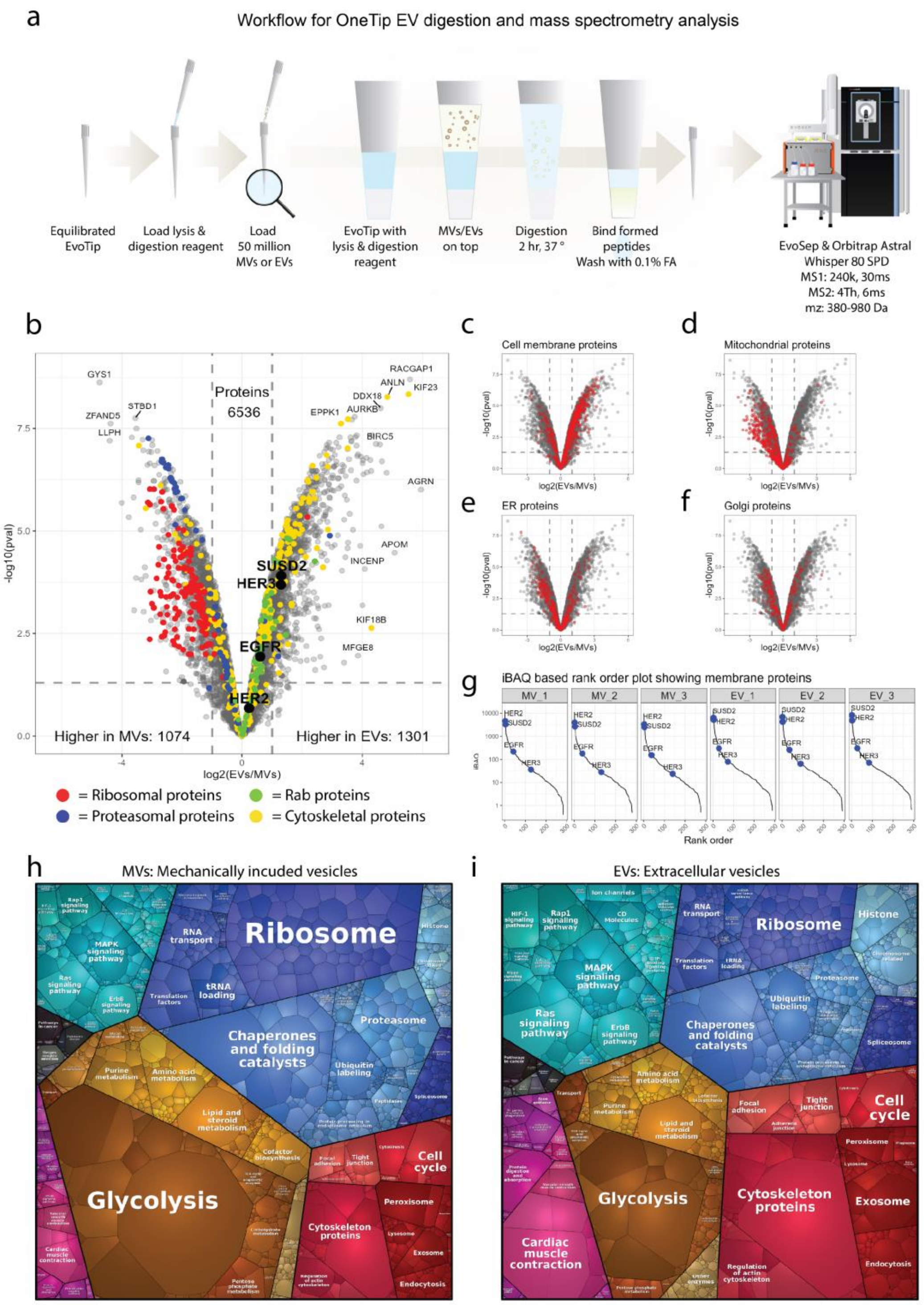


**Supplementary Figure 2. Mass spectrometry-based proteome analysis of mechanically induced vesicles (MVs) and extracellular vesicles (EVs) from SKBR3 cells. a.** Workflow for mass spectrometry-based analysis of MVs and EVs. **b.** Volcano plot showing proteins with different abundances in MVs and EVs; ribosomal proteins (red), proteasomal proteins (blue), Rab proteins (green), cytoskeletal proteins (yellow). **c-f.** Same volcano plot as in b with cell membrane proteins (c), mitochondrial proteins (d), endoplasmic reticulum proteins (e) and Golgi apparatus proteins (f) in red. **g:** Cell membrane proteins visualized in an iBAQ based rank order blot showing the abundance of HER2, SUSD2, EGFR and HER3 relative to all other cell membrane proteins. **h-i**. Proteome composition of MVs (h) and EVs (i) visualized in Voronoi plots. Each polygon represents the abundance of a quantified protein. The colors highlight different functional groups of proteins

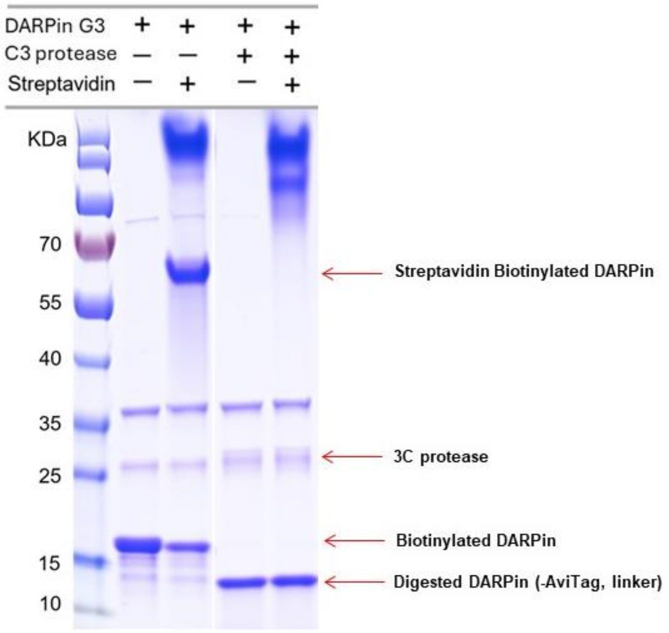


**Figure S3. Expression and purification of the biotinylated DARPins.** SDS-PAGE confirms the ability to attach DARPin to  streptavidin and release it by 3C protease cleavage.

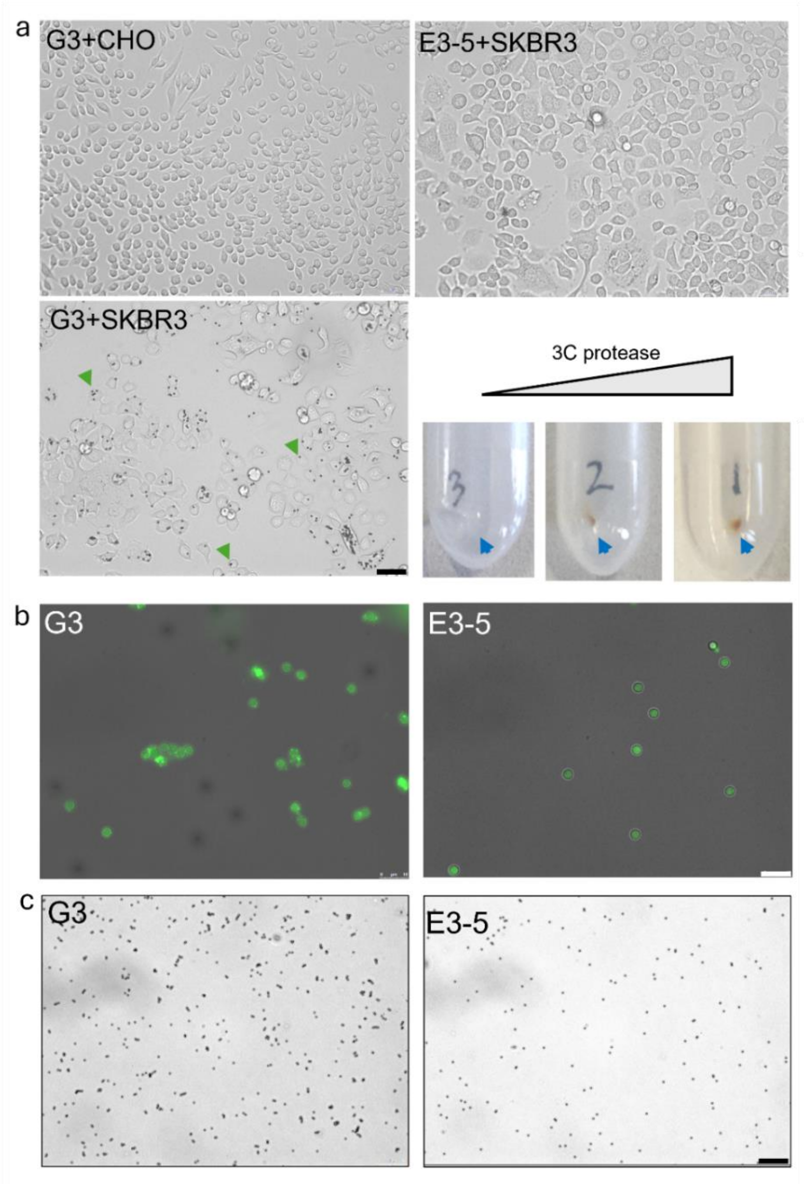


**Figure S4. The strategy used for purifying HER2 containing vesicles using different intact cells - calibration and controls.** a. Bright field light microscopy images showing CHO cells (HER2 negative, upper left), SKBR3 cells (HER2 positive, bottom left) incubated with magnetic beads (green arrows) conjugated with DARPin G3, and SKBR3 cells incubated with magnetic beads conjugated with DARPin E3-5 (not binding to HER2, control), upper right). The cells were washed with PBS prior to imaging to remove unbound magnetic beads. Magnetic beads are detected only after conjugation with DARPin G3 (Scale bar is 50 µm). The magnetic beads can be recovered by 3C protease followed by centrifugation, where increasing the amounts of 3C protease increase the recovery of magnetic beads (bottom right, blue arrows, 0 uM, 50 uM and 80 uM 3C protease were applied, respectively). **b.** Mechanically induced vesicles (MVs), produced from HER2-GFP expressing Sf9 cells, purified using magnetic force applied to magnetic beads conjugated with DARPins. The fluorescence images indicate the specificity of the process by the higher fluorescence from the HER2-GFP expressing Sf9 cells (Scale bar 10 µm). **c.** DARPin G3-coated magnetic beads show a tendency to aggregate after incubation with MVs from HER2-GFP expressing Sf9 cells, this was not observed with DARPin E3-5-coated beads (Scale bar 50 µm).

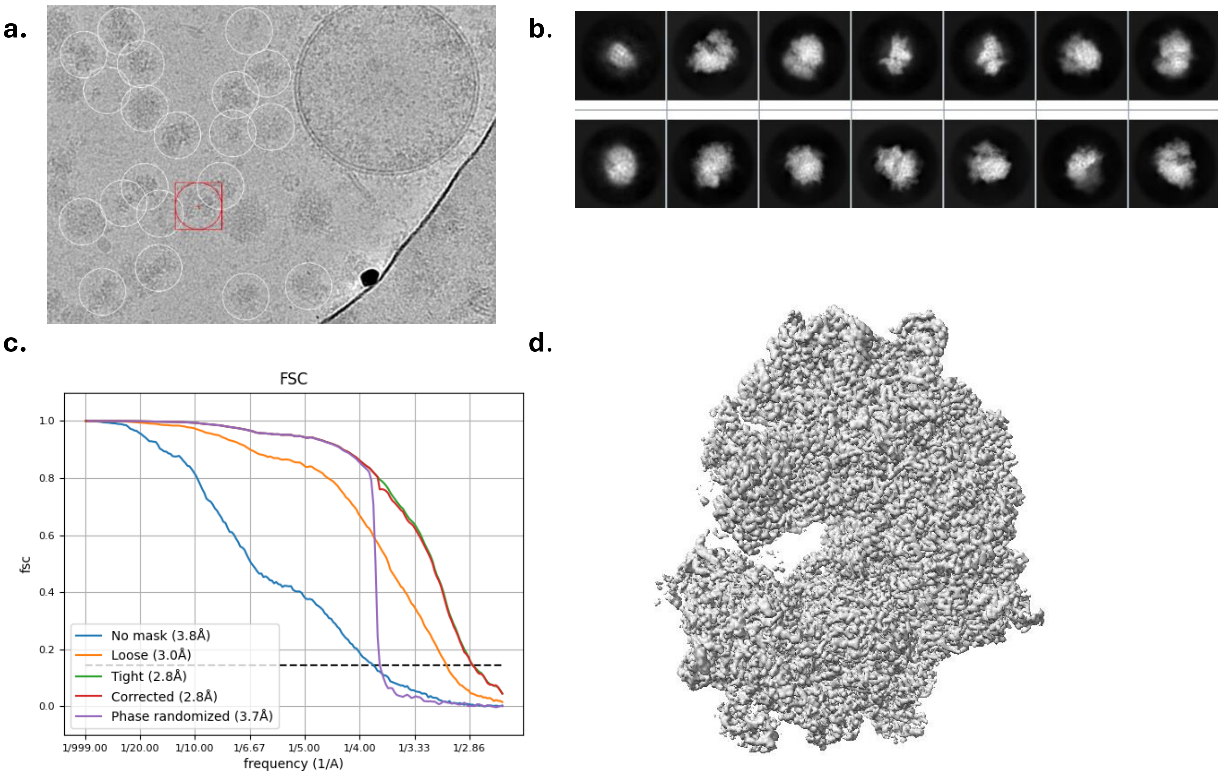


**Figure S5. The quality of the dataset was evaluated by structural characterization of the ribosome at high resolution. a.** Micrograph showing a vesicle and many ribosomes in the background (indicated using white circles). **b.** Reference-free 2D classes of ribosomes in HER2-vesicles. **c.** FSC gold-standard curves in the non-uniform refinement using cryoSparc. **d.** Final map after sharpening.

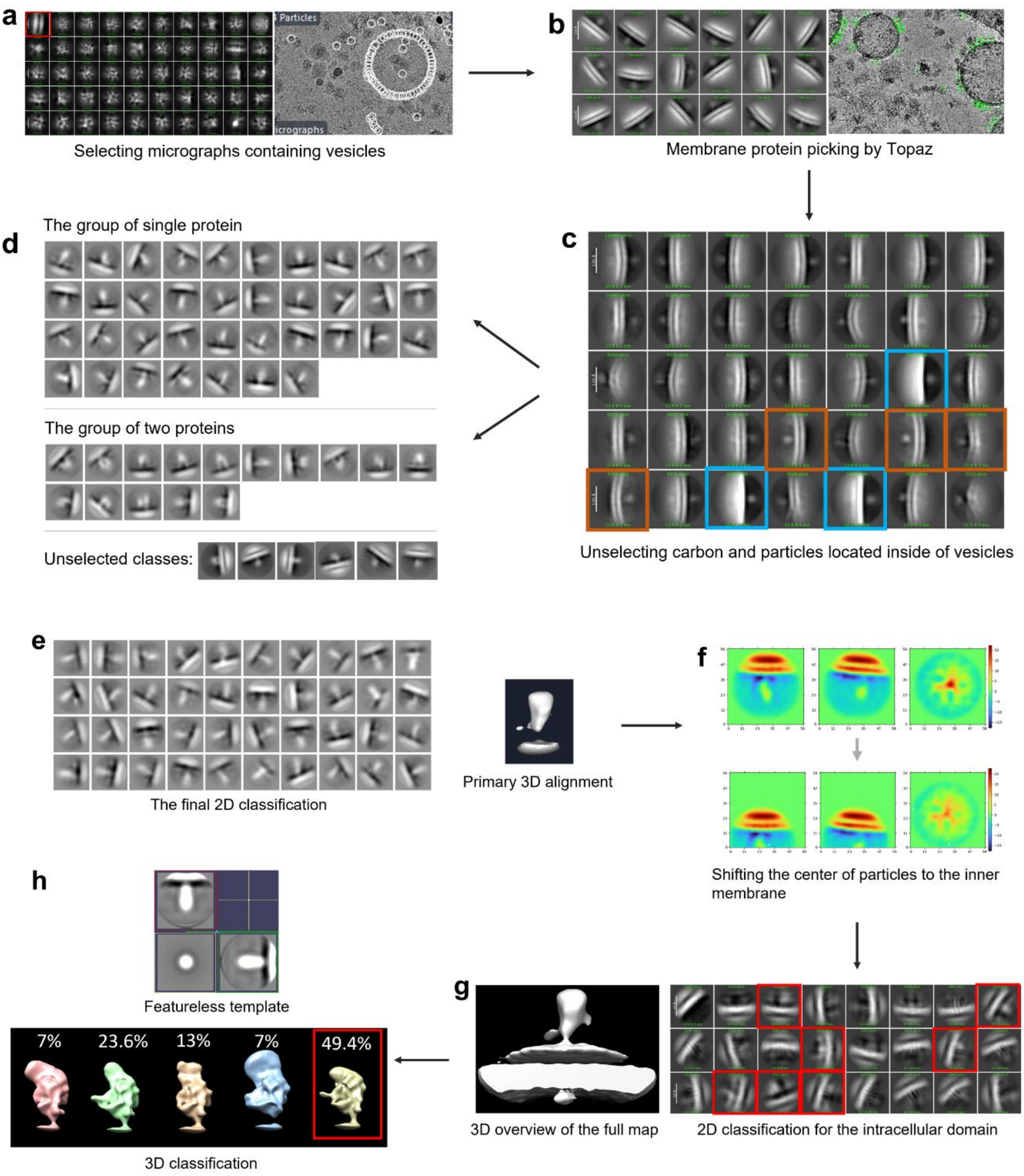


**Figure S6. Experimental workflow used for single particle cryo-EM analysis of receptors. a.** Selecting the micrographs containing vesicles and initial 2D averaging (left). **b.** Densities emanating from the membrane were picked by Topaz. **c.** Class averages which do not contain membranes were removed (Blue), in addition to concave membranes with respect to protein densities (red squares). **d.** Particle classification indicated a variety of protein architectures. The classes which show two proteins or proteins with non-relevant dimensions were removed. **e.** Resulting 2D classification of the dataset. **f.** The center of the images was shifted to the inner membrane to focus the analysis on the intracellular densities. **g.** Only classes that exhibit intracellular densities (the classes marked by the red squares) were selected for further structural determination. **h.** A 3D classification resulted in structures that resemble HER2, even though the template was a featureless structure (upper part, perpendicular views are shown).

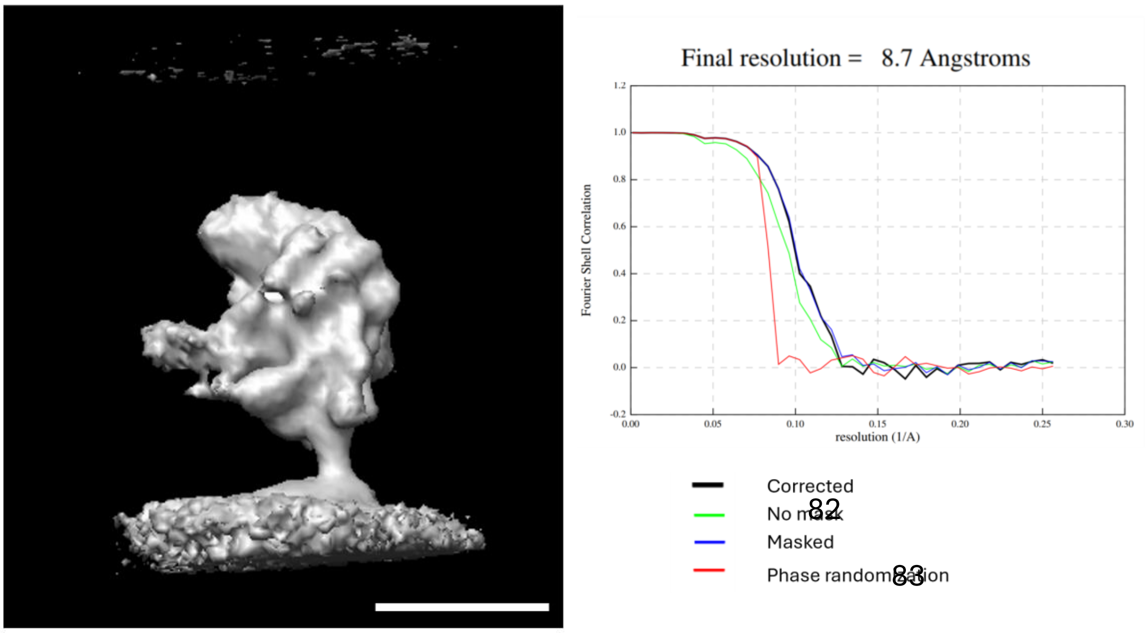


**Figure S7. The obtained structure of the HER2 receptor was resolved to 8.7 Å**. The rendered view of the resulting structure (left), resembling the HER2 structure, was refined to a resolution of 8.7 Å (right, gold-standard FCS resolution curves). The scale bar is 5 nm.

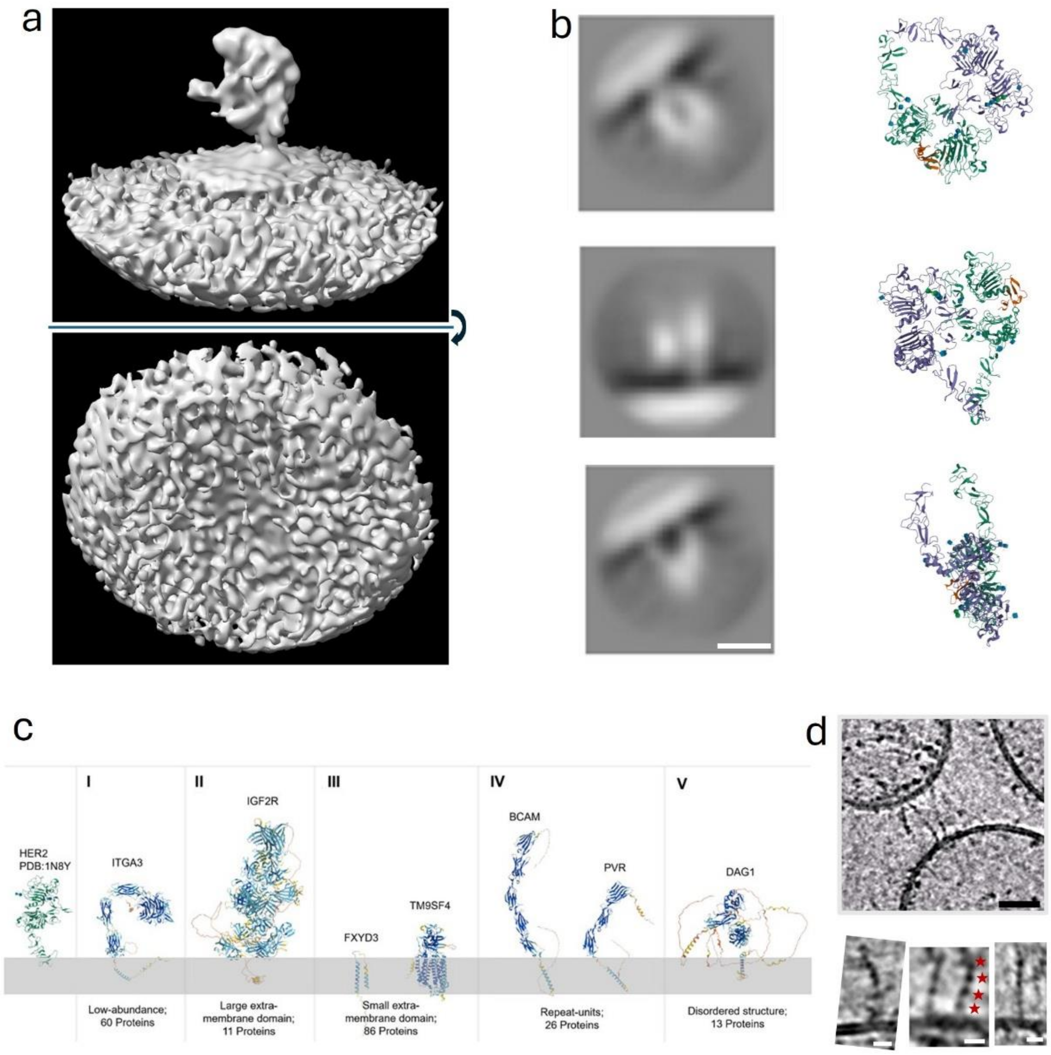


**Figure S8. Additional structural observations. a.** The refined structure does not exhibit an intracellular density, supporting the notion that this region is highly flexible in relation to the ECD. **b.** Representative 2D classes show a dimeric-like appearance in agreement with 2D views of the HER2-HER3 heterodimer (PDB 7MN5). Scale bar is 5 nm. **c**. AlphaFold2 models of the different classes of membrane proteins which were identified in MVs by mass spectrometry (Table 1) and compared with the structure of HER2 (Fig. 4c). **d**. Elongated membrane proteins composed of globular units span between vesicles (upper panel). Three examples from our data set is showed with higher magnification (lower panel). Repeating globular densities (red starts). Black scale bar 20 nm, white scale bar 10 nm.

**Table 1**. **Membrane proteins in diverse categories, correspond to Figure 4.** The gene names and UniProt IDs are shown. Cross correlation analysis between HER2 and the 34 protein with some structural resemblance are shown in Fig. 4c

**Membrane proteins with a large non-membrane domain (11 proteins)**

| Gene | Uniprot ID | Gene | Uniprot ID | Gene | Uniprot ID |
|---|---|---|---|---|---|
| IGF2R | P11717 | PTPRF | P10586 | PLXNB1 | O43157 |
| PLXNB2 | O15031 | SORL1 | Q92673 | NOTCH3 | Q9UM47 |
| GLG1 | Q92896 | L1CAM | P32004 | NUP210 | Q8TEM1 |
| PLXNA1 | Q9UIW2 | NOTCH2 | Q04721 | | |

**Membrane proteins with a small non-membrane domain (87 proteins)**

| Gene | Uniprot ID | Gene | Uniprot ID | Gene | Uniprot ID |
|---|---|---|---|---|---|
| FXYD3 | Q14802 | SLC35F6 | Q8N357 | LMAN2L | Q9H0V9 |
| TM9SF2 | Q99805 | TMED1 | Q13445 | TMX2 | G1K152 |
| TMED10 | P49755 | VTCN1 | Q7Z7D3 | SMIM7 | E9PP42 |
| LMAN2 | Q12907 | MPZL1 | O95297 | SDC4 | P31431 |
| LMAN1 | P49257 | TMX1 | Q9H3N1 | TLCD1 | Q96CP7 |
| TMED2 | Q15363 | JTB | O76095 | EMC10 | Q5UCC4 |
| CANX | P27824 | SPINT2 | O43291 | TMEM9 | B1ALM4 |
| TM9SF4 | Q92544 | GPR107 | Q5VW38 | GPR180 | Q86V85 |
| TMED4 | Q8R1V4 | CD99 | P14209 | ACP2 | P11117 |
| M6PR | P20645 | CCDC47 | J3KRX4 | TMX3 | Q96JJ7 |
| TACSTD2 | P09758 | GPR108 | Q9NPR9 | DDR1 | Q08345 |
| TMED7 | Q9Y3B3 | TMEM87A | Q8NBN3 | MFAP3L | D6RCF0 |
| TMEM165 | D6RBL0 | TM9SF1 | O15321 | F3 | Q86SE7 |
| TMED9 | Q9BVK6 | CD47 | Q08722 | SPPL2A | Q8TCT8 |
| RPN1 | P04843 | LAMP2 | P13473 | MPZL2 | O60487 |
| EPCAM | C9JKY3 | MLEC | F5H1S8 | KITLG | P21583 |
| LAMP1 | P11279 | TMEM194A | O14524 | TMCO3 | Q6UWJ1 |
| DHCR24 | Q15392 | PTTG1IP | P53801 | TMEM41A | Q96HV5 |
| TM9SF3 | Q9HD45 | SSR1 | P43307 | HLA-E | P13747 |
| SSR4 | P51571 | CD58 | P19256 | PGAP3 | Q96FM1 |
| SDC1 | P18827 | TMEM87B | Q96K49 | TPBG | Q13641 |
| TMED3 | Q9Y3Q3 | ATP6AP1 | Q15904 | TMEM9B | Q9NQ34 |
| TMEM123 | Q8N131 | MAGT1 | Q9H0U3 | ICOSLG | O75144 |
| DDOST | U3KQ84 | PROCR | Q9UNN8 | GLMP | Q8WWB7 |
| F11R | Q9Y624 | TGFBR1 | P36897 | UBAC2 | Q8NBM4 |
| PRSS8 | Q16651 | TNFRSF12A | Q9NP84 | HLA-C | P10321 |
| CA12 | O43570 | ATP6AP2 | O75787 | SIDT2 | Q8NBJ9 |
| TMEM109 | Q9BVC6 | TUSC3 | Q13454 | SLC39A14 | Q75T53 |
| TMED5 | Q9Y3A6 | PIGK | Q92643 | DCD | P81605 |

**Membrane proteins with non-membrane repeating domains (26 proteins)**

| Gene | Uniprot ID | Gene | Uniprot ID | Gene | Uniprot ID |
|---|---|---|---|---|---|
| BCAM | P50895 | NOMO3 | J3KN36 | DSC2 | Q02487 |
| PTGFRN | Q9P2B2 | DSG2 | Q14126 | PTK7 | Q13308 |
| PROM2 | Q8N271 | ALCAM | Q13740 | PCDH1 | Q08174 |
| CD97 | P48960 | ITGA2 | P17301 | BSG | P35613 |
| IGSF3 | O75054 | PTPRK | Q15262 | PVR | P15151 |
| CD46 | P15529 | PVRL4 | Q96NY8 | CADM1 | Q9BY67 |
| EPHB3 | P54753 | PVRL2 | Q92692 | EPHA1 | P21709 |
| NPTN | Q9Y639 | PVRL1 | Q15223 | CD276 | Q5ZPR3 |
| IGSF8 | Q969P0 | EPHA2 | P29317 | | |

**Membrane proteins with intrinsically disordered non-membrane domains (13 proteins)**

| Gene | Uniprot ID | Gene | Uniprot ID | Gene | Uniprot ID |
|---|---|---|---|---|---|
| TGOLN2 | O43493 | SLC39A10 | Q9ULF5 | SYVN1 | Q86TM6 |
| DAG1 | Q14118 | SLC39A6 | Q13433 | MMP15 | P51511 |
| VASN | Q6EMK4 | MIA3 | Q5JRA6 | CD44 | P16070 |
| CLCC1 | Q96S66 | STIM1 | Q13586 | | |
| APP | P05067 | APLP2 | Q06481 | | |

**Membrane proteins with some resemblance to HER2 (34 proteins)**

| Gene | Uniprot ID | Gene | Uniprot ID | Gene | Uniprot ID |
|---|---|---|---|---|---|
| HER2 | P04626 | ITGB1 | P05556 | PTPRA | P18433 |
| SUSD2 | Q9UGT4 | NCSTN | Q92542 | ENG | P17813 |
| RPN2 | Q3SZI6 | GPR56 | A0A024R6U7 | NCLN | B2RA56 |
| SORT1 | Q99523 | KIAA1324 | A0A6P5P8N1 | ADAM9 | Q13443 |
| EGFR | P00533 | ITGB6 | P18564 | QSOX1 | O00391 |
| ADAM10 | O14672 | PIGT | Q969N2 | LDLR | P01130 |
| ALPP | P05187 | QSOX2 | Q6ZRP7 | ADAM17 | P78536 |
| TAPBP | O15533 | SEL1L | Q9UBV2 | EMC1 | Q8N766 |
| ITGAV | P06756 | KIAA2013 | Q8IYS2 | ATRN | O75882 |
| ITGB5 | P18084 | TMEM132A | Q24JP5 | NPC1 | A0A193DRS0 |
| CPD | O75976 | ADAM15 | Q13444 | | |
| ITFG1 | Q8TB96 | ERBB3 | P21860 | | |
